# Direct excitatory synapses between neurons and tumor cells drive brain metastatic seeding of breast cancer and melanoma

**DOI:** 10.1101/2024.01.08.574608

**Authors:** V. Venkataramani, M.A. Karreman, L.C. Nguyen, C. Tehranian, N. Hebach, C.D. Mayer, L. Meyer, S.S. Mughal, G. Reifenberger, J. Felsberg, K. Köhrer, M.C. Schubert, D. Westphal, M.O. Breckwoldt, B. Brors, W. Wick, T. Kuner, F. Winkler

## Abstract

Interactions between neurons and cancer cells are found in many malignancies, but their relevance for metastatic organ colonization remain largely unknown. It is also unclear whether any direct synaptic communication between neurons and cancer cells of non-neural tumor types exists, and if so, whether this can support metastasis and thus cancer progression. Here we show that excitatory synapses are formed between neurons and brain-metastatic melanoma and breast cancer cells. This starts at an early microscopic stage after extravasation into the brain parenchyma, during residence of cancer cells in the perivascular niche, a critical step for their survival. These neuron-cancer synapses showed a *bona fide* synaptic ultrastructure, and generated excitatory postsynaptic currents mediated by glutamate receptors of the AMPA subtype in cancer cells. In accordance, AMPA receptor signatures were consistently detected in preclinical and patient samples of melanoma and breast cancer brain metastases. Genetic perturbation and pharmacological inhibition of AMPA receptors with the approved antiepileptic drug perampanel in models of breast and melanoma cancer reduced the number of brain metastases and overall brain metastatic burden. These findings demonstrate for the first time that neurons can form biologically relevant direct synapses with non-neural cancer cells. In brain metastasis, a particularly challenging complication of many common malignancies, this non-canonical stimulatory synaptic interaction offers novel therapeutic opportunities.

## Introduction

The nervous system can contribute to the development and progression of both intra- and extracranial tumors through several mechanisms^1, 2^. One of the most unexpected findings was the formation of excitatory synapses between presynaptic neurons and postsynaptic cancer cells, which has been reported to stimulate tumor growth and invasion in certain cancer types of neural origin^3–6^. While the finding of such neuron-cancer synapses stimulated the emerging field of cancer neuroscience, it remained unclear whether they are simply shadows of their neural progeny. Oligodendrocyte precursor cells (OPCs), the neural putative cell of origin of glioma types^7^ that receive glutamatergic synaptic input^3–5^, receive such input in their nonmalignant physiological state^8^. In mammals, *bona-fide* synapses between neurons and non-neuronal cells are limited to two targets: next to OPCs, only muscle cells are known to receive synaptic input^9^. Yet, neuromuscular synapses contain a basal membrane that separates the pre- and postsynaptic compartments. It is an exciting question whether neurons can form direct synapses with other cell types, too, including cancer cells from non-neural origin.

Brain metastases (BM) are associated with high morbidity and mortality and occur in as many as 10%- 40% of all cancer patients^10^, with increasing incidence and dismal prognosis^11^, most often originating from breast cancer (triple negative or Her2 positive), lung cancer, or melanoma. There is an unmet clinical need for novel therapeutic approaches to treat or preferably prevent this devastating disease^11^. In breast cancer brain metastases, cancer cells can reside next to a synapse between two neurons, with glutamate spilled over from the synaptic cleft stimulating NMDA receptors (NDMARs) which in turn fosters the growth of established macrometastases^12^. However, these indirect contacts rather resemble those with astrocytes, and effective pharmacological targeting of NMDARs is generally challenging because of intolerable side effects that occur even with mild levels of inhibition^13^. Thus, the question remained whether functional synapses between neurons and brain-invasive cells of various non-neural cancer types can also be found, and if so, at which step of the metastatic cascade^14^, how this influences early organ colonization, and whether the synapse type is more amenable to pharmacological inhibition.

## Results

### Neuronal activity-related cancer cell Ca^2+^ transients in a crucial metastatic niche

When circulating breast cancer and melanoma cells from various tumor models left the blood vessel and extravasated into the mouse brain, they were consistently found in a perivascular niche (**Fig. 1a-c**). Intravital time-lapse multiphoton microscopy following the long-term fate of these cells revealed that this striking perivascular position was strictly maintained by cancer cells even after cell divisions and during early growth (**Fig. 1b,c**), in line with previous findings^14^. This makes strict perpetuation of the perivascular niche position to a mandatory step of the brain metastatic cascade and suggests a survival-promoting function of this niche for non-neural, metastatic cancer cells. During metastatic seeding of the perivascular niche and early proliferation therein, we observed Ca^2+^ transients *in vivo* (**Fig. 1d-e, Supplementary Videos 1 and 2**). The fraction of metastatic lesions exhibiting such Ca^2+^ transients increased over time (**Fig. 1f**). Ca^2+^ activity in breast cancer micro-metastases coincided with increased growth when compared to Ca^2+^-silent BM (**Fig. 1g**). Neuronal stimulation can result in such increases in cytosolic Ca^2+^ concentration, either via influx from extracellular space or release from intracellular stores^3, 15^. Importantly, previous studies in glioma^3–5^ established a close link between glutamatergic neuron-cancer synaptic stimulation and Ca^2+^ transients in cancer cells. To test the potential impact of neuronal activity on metastatic cancer cells, intravital microscopy in awake mice bearing breast cancer or melanoma metastases was performed (**Fig. 1h-j**; top panels) and compared to ketamine/xylazine anesthesia conditions (**Fig. 1h-j** bottom panels). Ketamine/xylazine anesthesia leads to strongly reduced overall neuronal activity and affects both NMDARs and AMPARs^16, 17^, but should not affect potential tumor cell-autonomous generation of Ca^2+^ transients and their potential exchange via intercellular gap junctions^18^. A significant reduction of cancer cell Ca^2+^ transients was evident in mice anesthetized with ketamine/xylazin, compared to the Ca^2+^ transients recorded in the same brain regions when these mice were awake (**Fig. 1k, Supplementary Video 3**). Together, this was the first indication for the existence of a functional communication between neurons and metastatic cancer cells in the brain beginning at crucial steps of early brain seeding, with potential influence on subsequent BM growth.

**Figure 1:**
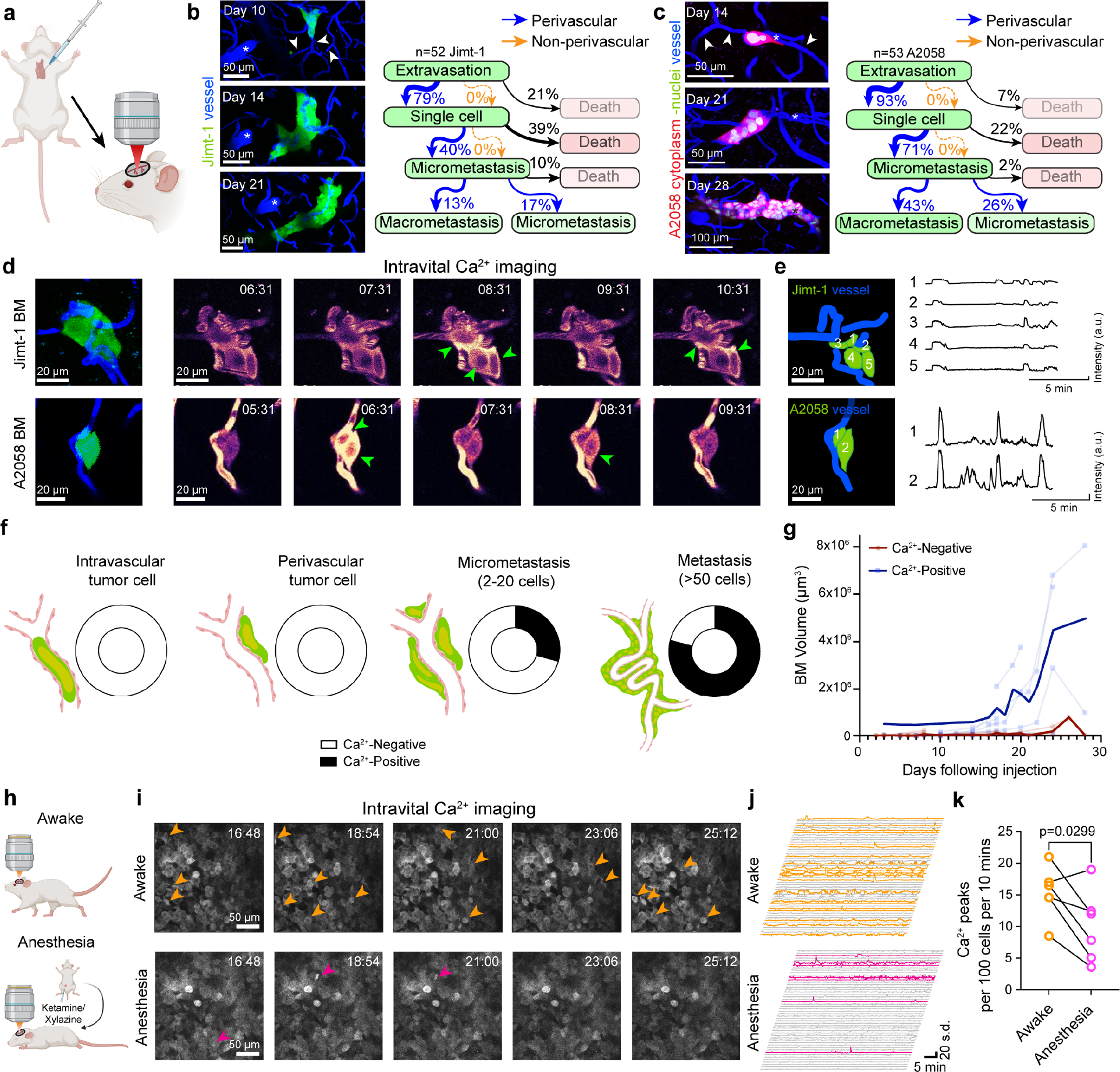
Ca^2+^ activity in early stage breast cancer and melanoma brain metastatic cells *in vivo* **a**, Experimental scheme of *in vivo* intravital microscopy of Jimt-1 breast cancer and A2058 melanoma BM following intracardial injection. **b,c,** Long-term intravital microscopy of breast cancer (Jimt-1) and melanoma (A2058) brain-seeding metastatic cancer cells. In the top panels, at the micro-metastatic stage, the arrowheads depict the vessels that are later used for perivascular metastatic growth. Quantification of the metastatic cascade demonstrates exclusive survival and proliferation in the perivascular niche. **d**, Left panels: intravital microscopy of Jimt-1 (top) and A2058 (bottom). Tumor cells shown in green, vessel lumen shown in blue. Right: Representative time series recordings of Ca^2+^ activity in micro-metastases shown in left panels, expressing the fluorescent Ca^2+^ sensors. Intravascular injected TRITC dye is visible in the same channel. Arrowheads indicated fluctuations in intacellular Ca^2+^ concentration. Time is given as min:s. **e,** Left panels: segmentation of the tumor cells (green, numbered) and capillaries (blue) of the micro-metastases shown in d. Ca^2+^ traces for each numbered tumor cell in the field of view are shown on the right (see Supplementary videos 1 and 2). **f**, Quantification of Ca^2+^ activity during various stages of the brain metastatic cascade; arrested, intravascular tumor cell (n=8); single perivascular tumor cell (n=9); early micrometastasis (n=17) consisting of few tumor cells and multi-cellular metastasis (n=24). Ca^2+^ imaging datasets were obtained from N=6 mice. **g,** Quantification of *in vivo* growth of BM that do not show Ca^2+^ activity (n=26) and of those BM that do (n=15), monitored by intravital microscopy following intracardiac injection. **h,** Experimental scheme of *in vivo* Ca^2+^ imaging of A2058 melanoma cells in awake (top) or ketamine/xylazine anesthetized (bottom) mice after intracranial tumor injection. Same BM regions were recorded while the mouse was awake and moving freely or while under ketamine/xylazine-induced anesthesia. **i,** Representative time series recordings of Ca^2+^ transients in A2058 melanoma brain tumor cells *in vivo* in awake or anesthetized mice. Arrowheads indicate intracellular Ca^2+^ fluctuations. **j**, Ca^2+^ traces from representative A2058 MBM cells in awake and anesthetized mice. Traces in orange (awake) and magenta (anesthetized) are traces from cells showing Ca^2+^ transients during the period of recording. **k,** Number of Ca^2+^ peaks per 100 A2058 MBM cells in 10 min of *in vivo* imaging in awake vs. anesthetized mice (n=6 regions, 465-1244 cells per region, from N=3 mice). Normality was confirmed with a Shapiro-Wilk test, p value was determined with a two-tailed ratio paired T-test.

### Synapses between neurons and brain metastatic cancer cells generate AMPA receptor-mediated postsynaptic currents

To further investigate if synaptic contacts between neurons and brain metastatic cells exist, we performed whole-cell patch-clamp recordings of single breast cancer and melanoma cells in acute brain slices of mice during early growth in the perivascular niche after extravasation, and on co-cultures of brain metastatic cancer cells, neurons and astrocytes (**Fig. 2a,b**; **Supplementary** Fig. 1a). In all conditions, our recordings showed spontaneous excitatory postsynaptic currents (sEPSCs) in a subset of tumor cells from the brain metastatic breast cancer and melanoma cell lines, Jimt-1, E0771 and A2058 (**Fig. 2c**), demonstrating functional synapses between presynaptic neurons and postsynaptic tumor cells. The observed currents showed a fast rise time and exponential decay, as hallmark features of AMPA receptor-mediated sEPSCs (**Fig. 2d**, **Supplementary** Fig. 1b-f). sEPSCs in both breast cancer- and melanoma brain metastatic cells were completely blocked by the AMPAR-specific antagonist cyanquixaline (CNQX), confirming that AMPARs are functionally contributing to synapses between neurons and metastatic cancer cells (**Fig. 2e**). Aside from sEPSCs, slow inward currents (SIC) were also observed, but only in co-cultures of neurons with brain-tropic melanoma cells, and only in one cell line (**Fig 2f-h**). Importantly, voltage-activated currents were found (lower trace in **Fig. 2i**) while action potentials could not be elicited in brain metastatic breast cancer and melanoma cells, suggesting that these cells cannot generate regenerative potentials (**Fig. 2i**). Rather, they are the receivers of unidirectional synaptic input from neurons. Taken together, these data demonstrate excitatory postsynaptic currents, mediated through synapses between neurons and individual metastatic cancer cells ("neuron-BM synapses", NBMS) containing glutamate receptors of the AMPA subtype, both in brain metastatic breast cancer and melanoma cells, with cancer cells on the postsynaptic (receiving) side of this electrochemical communication path.

**Figure 2:**
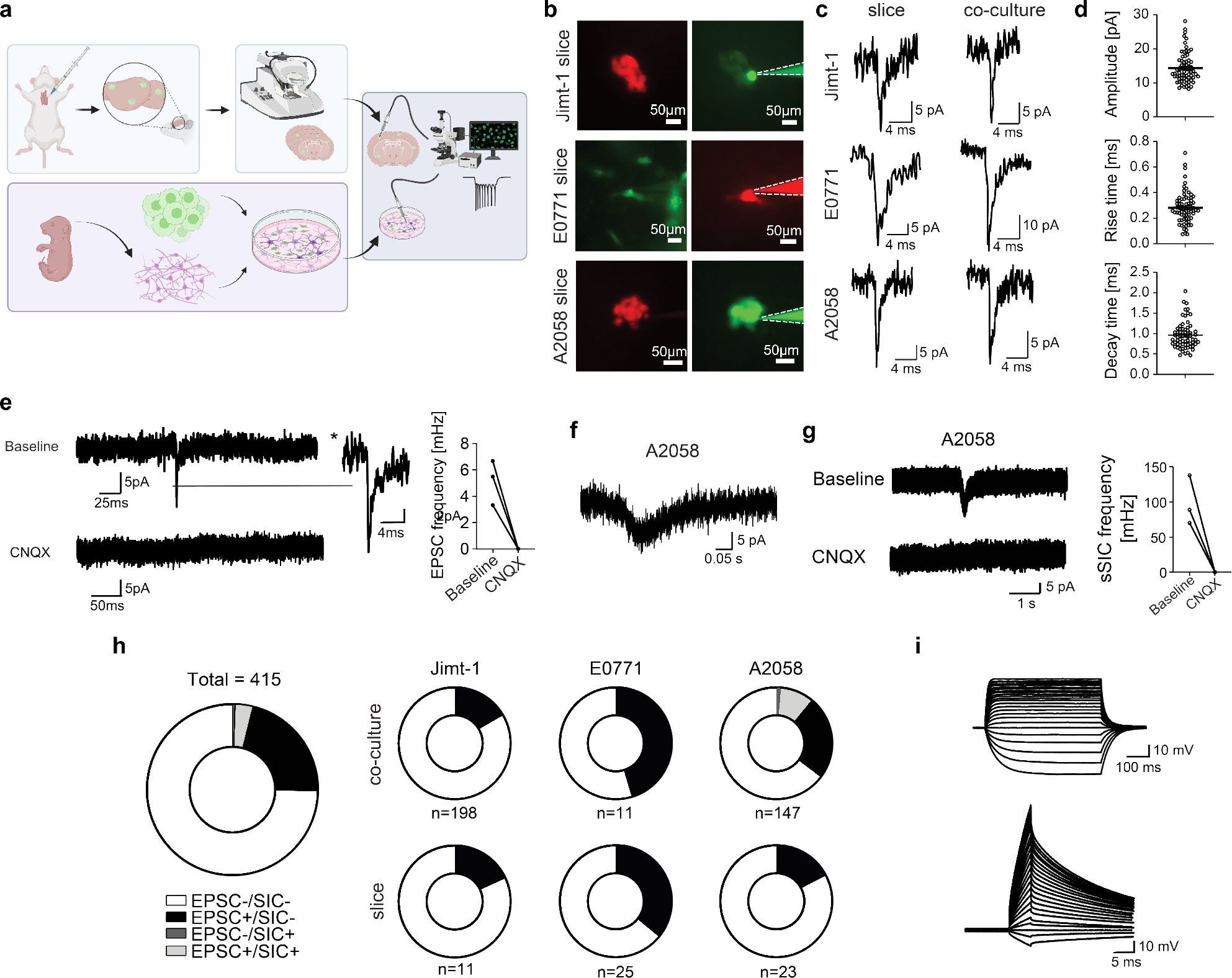
Functional characterisation of neuron-BM synapses **a**, Workflow for whole-cell patch-clamp electrophysiology *ex vivo* and *in vitro*. BM cells are either injected intracardially in mice or were co-cultured with neurons before whole-cell patch clamp experiments. **b**, Representative images of whole-cell recordings from micrometastases in acute brain slices (Jimt-1 expressing pan-cellular tdTomato, red; E0771 (mouse mammary cancer): pan-cellular GFP, green, A2058 expressing cytoplasmic tdTomato and nuclear GFP) and filled with a fluorescent dye (either Alexa 488 or Alexa 594) via the patchpipette to verify the identity of the cell recorded from (right panels). **c,** Representative sEPSCs recorded in different BM cell lines in acute brain slices (left) or in co-culture with neurons (right). **d,** Quantitative properties of sEPSCs in Jimt-1 co-cultures indicative of AMPA receptor kinetics. **e,** Traces showing inhibition of sEPSCs following application of CNQX in Jimt-1 co-culture and corresponding quantification. **f**,**g**, Representative traces of A2058 melanoma BM cells showing spontaneous SICs (g), and their inhibition by CNQX. Traces before and after CNQX are shown in g, and their quantification (n=3). **h**, Quantification of different electrophysiological subgroups of BM cells (Jimt-1, E0771 and A2058) in vitro (co-culture) and ex vivo (acute brain slices). **i**. Representative current-clamp recordings from a Jimt-1 BM cell. Voltage responses to current injections are shown (n=11 cells, n=29 cells).

### Ultrastructural confirmation of neuron-BM synapses

Next, we assessed the ultrastructural correlate of a glutamatergic synapse formed between neurons and BM cells. To establish whether and at which stage during the brain metastatic cascade^14^ NBMS are formed, electron microscopy (EM) of preclinical BM at different time points after intracardial injection of breast cancer and melanoma cells was performed. To be able to identify the earliest stages of brain seeding and to relocate these cells for subsequent EM, we performed intravital correlative microscopy^19, 20^ to identify brain-colonizing cancer cells using *in vivo* microscopy through a cranial window in mice, followed by three-dimensional EM (3DEM) of these exact regions (**Supplementary** Fig. 2a).

Earliest BM (<5 cells per metastatic lesion) were successfully targeted, and clear NBMS were detectable (**Fig. 3a, b**, **Supplementary** Fig. 2b, **Supplementary Video 4**) in 20% of single breast cancer cells or micrometastases in the perivascular niche (**Fig. 3c**). Furthermore, serial section scanning electron microscopy^21, 22^ from two models of melanoma BM also revealed NBMS in the micrometastatic stage (**Fig. 3d-f**, **Supplementary** Fig. 2c, **Supplementary Video 5-6**). Synaptic contacts formed onto brain metastatic cells were counted as such if 2 out of 3 criteria were met: (1) synaptic vesicle cluster present in neuronal presynaptic compartment, (2) synaptic cleft visible, (3) postsynaptic density (PSD) apparent in tumor cells (**Fig. 3a, d**, right panels). Importantly, the cancer cell always harbored the post-synapse and never showed pre-synaptic features in accordance with the functional electrophysiology data (**Fig. 2**; **Fig. 3a**, **b, d** and **e**). Further analysis revealed that the vast majority of NBMS were indeed direct synapses between presynaptic neurons and postsynaptic cancer cells, without a non-malignant neuronal structure co-located on the postsynaptic side (**Fig. 3**). The latter speaks against a frequent hijacking of pre-existing brain synapses by cancer cells, in contrast to findings in glioma^3^, implying frequent *de novo* synaptogenesis in brain metastasis.

**Figure 3:**
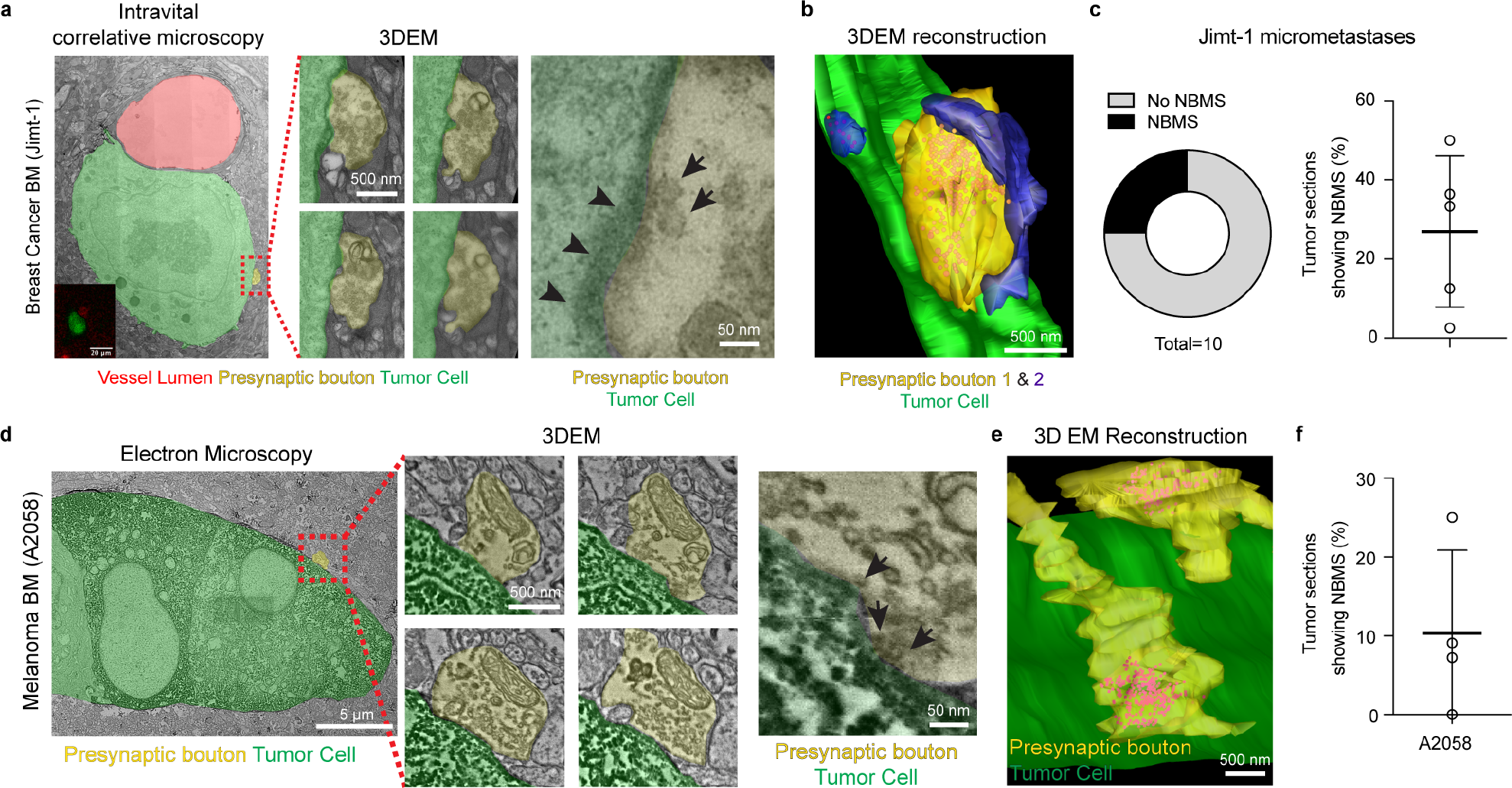
Ultrastructure of neuron-BM synapses in breast cancer and melanoma. **a**, Intravital correlative 3D electron microscopy (3DEM) shows a neuronal presynaptic bouton (yellow) on a postsynaptic Jimt-1 single perivascular cancer cell (4 days post-intracardiac injection, green). Arrowheads show postsynaptic cleft and arrows indicate synaptic vesicles. **b**, 3D EM reconstruction of the tumor-brain metastasis synapse shown in a, synaptic vesicles shown in red. **c**, Quantification of NBMS on n=10 Jimt-1 micrometastases (3-7 days post-intracardiac injection, n= 5 mice), and the percentage of randomly selected tumor cell sections showing NBMS (80 sections from n=5 tissue blocks from 3 mice). **d**, Immuno-electron microscopy of DAB (3,3′-Diaminobenzidine)-marked A2058 melanoma brain micrometastasis, DAB can be recognised as electron-dense deposits in the tumor cell cytoplasm. Melanoma BM is marked in green, presynaptic bouton in yellow. Arrows indicate synaptic vesicles. **e**, 3D EM reconstructions show the presynaptic vesicles (red). **f**, Quantification of NBMS on n=4 A2058 micrometastases (14 days post-intracardiac injection), and the percentage of randomly selected tumor cell sections (n=96) showing NBMS (n=4 tissue blocks from 3 mice).

### Molecular characterization of neuron-BM synapses

Next, we set out to identify the molecular composition of NBMS in preclinical models and clinical samples. First, a dataset was analyzed in which mRNA sequencing was performed of cancer cells isolated from various preclinical models (14-21 days following forebrain injection) compared to the same cancer cells in 2D culture^23^ (**Fig. 4a**). Expression of AMPAR subunit-genes *GRIA2, GRIA3* and *GRIA4* were upregulated in breast and lung cancer and melanoma cells growing in the brain, while differences in expression of genes associated with other neurotransmitter receptors were less consistent and less pronounced (**Fig. 4b**, **Supplementary** Fig. 3a). We then analyzed RNA sequencing data from the PreventBM cohort of 95 patients with brain metastases from different primary cancers, and first determined the expression of the AMPAR subunit genes. *GRIA1-4* were found to be expressed across BMs from melanoma and breast cancer; interestingly, this appears to be a general feature across cancer entities since lung-, colon- and kidney cancer BM also expressed *GRIA1-4* (**Fig. 4c**). To assess if the expression of the AMPAR subunits is enriched in BM, and how this compares to the primary tumor, we interrogated gene expression data from primary breast cancers, lung adenocarcinomas and melanomas from The Cancer Genome Atlas (TCGA). *GRIA1-GRIA4* were generally higher expressed in BM as compared to their corresponding primaries (**Fig 4d**,**e**, **Supplementary** Fig. 3b-e). The “AMPAR score”, defined as the sum of the z-scores over *GRIA1-4* mRNA expression (**Supplementary** Fig. 3e), was at a similar level across all entities (**Fig 4f**, **Supplementary** Fig. 3f), supporting a general ability of cancers to participate in AMPAR synaptic signalling. Comparing the AMPAR score between BM and primary tumor samples in the PreventBM cohort, and in primary tumor samples from TCGA, a consistently higher AMPAR score was found in BM samples compared to primary tumors (**Fig. 4f**). In the non-pathological brain, the GRIA2 subunit is RNA-edited on the Q/R site. Homomeric and heteromeric AMPAR containing the edited form GRIA2R show low Ca^2+^ conductance^3, 24^, whereas the underedited GRIA2Q subunit renders AMPAR Ca^2+^-permeable. Thus, we next set out to investigate the editing status of *GRIA2* expressed in BM samples. From 95 BM patients, the editing status could be technically assessed from 14, of which 71% (10/14) showed a clear reduction of editing of *GRIA2* (**Fig 4g**). Since the editing status of GRIA2 is close to 100% in the adult normal brain, this finding indicates that GRIA2 expression is derived from neoplastic cells, at least partially, and not the brain microenvironment. Indeed, interrogation of a single cell RNA-sequencing dataset from paired human melanoma BM and extracranial metastases revealed that AMPAR genes are higher expressed in BM tumor cells as compared to patient-matched extracranial melanoma metastases^25^ (**Fig. 4h**). Collectively, these findings provide molecular evidence for AMPA receptor expression in human brain metastases from multiple tumor entities, its upregulation in the brain environment during metastatic growth, and an explanation for AMPAR-mediated Ca^2+^ signals in BM cells.

**Figure 4:**
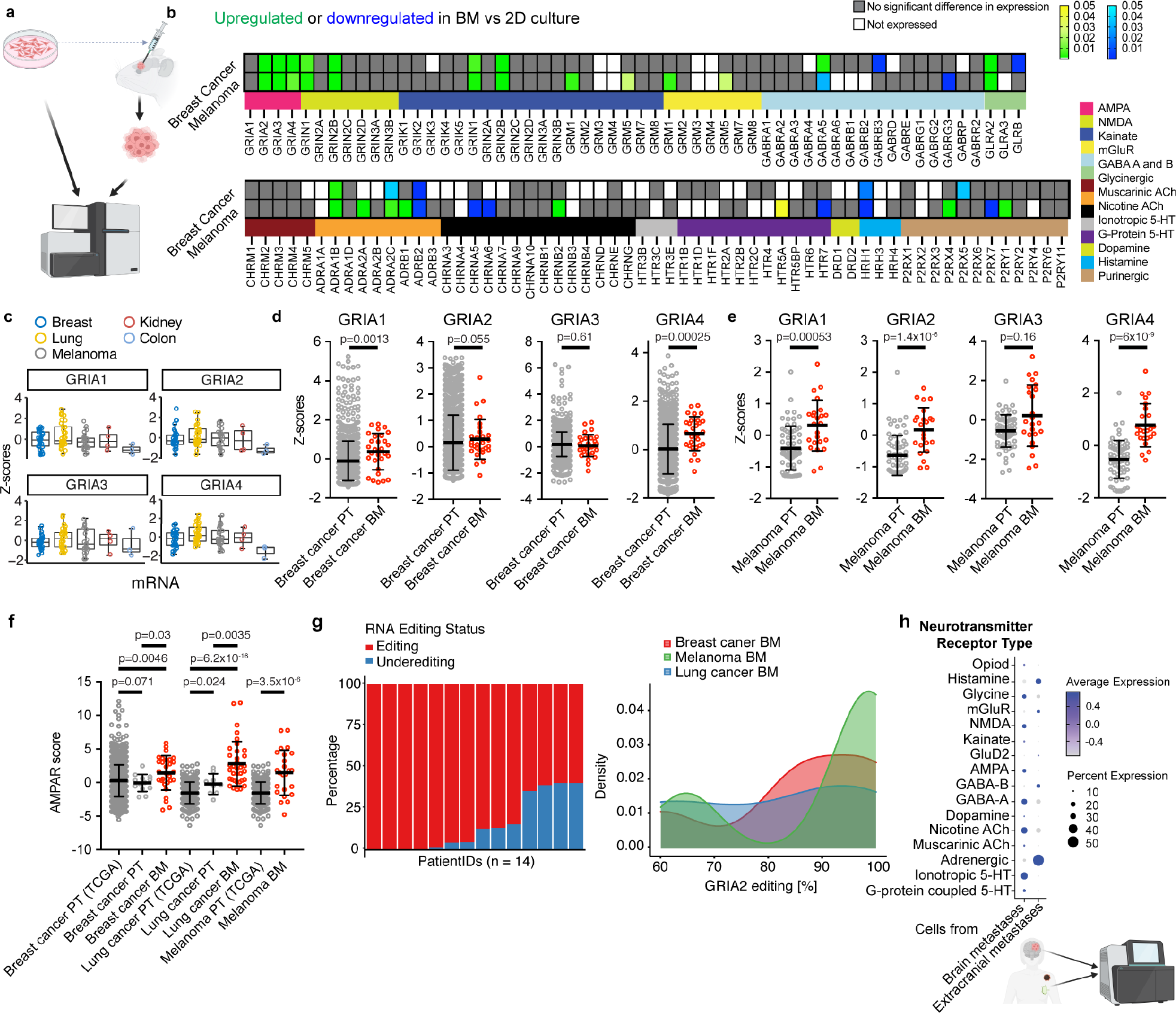
Molecular synaptic signatures in primary tumors, brain and extracranial metastases. **a**, Schematic summary of the workflow comparing RNA expression in human breast cancer and melanoma cells lines grown in the mouse brain or in 2D cell culture. Based on Wingrove *et al*^18^. **b,** Changed regulation of neurotransmitter receptor genes (indicated below) in human melanoma BM (A375- Br) and breast cancer BM (MDA-MB-231-Br2) cells grown in 2D-culture vs. in the forebrain of mice^18^. Significant upregulation is indicated in yellow/green, downregulation in blue. Grey indicates no significant difference in RNA expression. AMPAR genes *GRIA2-4* are significantly upregulated in melanoma and breast cancer BM cells grown in the brain compared to 2D culture. **c**, RNA sequencing of patient brain metastases from different entities (n=29 from breast cancer BM, n=13 from breast cancer PT; n=35 from lung cancer BM, n=9 from lung cancer PT, n=24 from melanoma, n=4 from kidney cancer, and n=3 from colon cancer). Expression of AMPAR genes *GRIA1-4* in brain metastases from different entities. **d,e**, Expression of AMPAR genes *GRIA1-4* in breast cancer (**d**, n=29) or melanoma (**e**, n=24) BM vs primary tumors (PT, 1102 breast cancers and 103 melanomas from The Cancer Genome Atlas (TCGA)). **f**, Comparison of AMPAR scores between BM and primary tumor (PT) datasets. BC: breast cancer, LUAD: lung adenocarcinoma; Mel: Melanoma; BM: brain metastases; PT: primary tumor. Medians are shown, and error bars represent standard deviation. P-values were determined with a two-sided Wilcoxon test. **g**, Stacked barplot (left) displaying the proportion of *GRIA2* mRNA editing (in %) in 14 BM patients. Right, density plot for GRIA2 mRNA editing in patients with BM from different entities. **h**, Single cell RNA sequencing of extracranial and matched intracranial metastases^20^ of melanoma show expression of neurotransmitter receptor genes. AMPA genes are higher expressed in brain metastases versus extracranial metastases.

### Genetic and pharmacological perturbation of AMPAR inhibits brain metastatic outgrowth

Finally, we sought to investigate how the functionality of NBMS influences the outgrowth of BM in preclinical models. Hereto, brain-tropic tdTomato expressing Jimt-1 breast cancer cells were transduced with a dominant negative AMPAR subunit fused with GFP (GluA2-DN-GFP), or the GFP-tagged WT AMPAR subunit GluA2. To assess the effect of AMPAR functionality on BM growth, mice were intracardially injected with GluA2-DN-GFP or GluA2-GFP cell lines and sacrificed 28 days post-injection. Mice injected with GluA2-DN-GFP expressing Jimt-1 showed a significantly lower number of BM, resulting in overall lower metastatic burden compared to mice injected with Jimt-1 GluA2-GFP (**Fig 5a**, **b**). To further confirm the role of AMPAR on BM growth *in vivo*, and, importantly, to probe a potential road for clinical translation, we set out to inhibit AMPAR function pharmacologically with perampanel. Perampanel is a selective and non-competitive AMPAR antagonist and is an FDA-approved drug for the treatment of epilepsy^26^. Administration of perampanel resulted in a lower metastatic burden, fewer BM per mouse, and a lower volume per BM in mice that were intracardially injected with breast cancer cells (**Fig 5c**, **d**). In line, in a melanoma BM model, perampanel treatment also reduced BM growth *in vivo* (**Fig 5e**). Taken together, these findings demonstrate that inhibition of glutamatergic NBMS reduces BM growth in pre- clinical models and suggests a therapeutic window for clinical translation.

**Figure 5:**
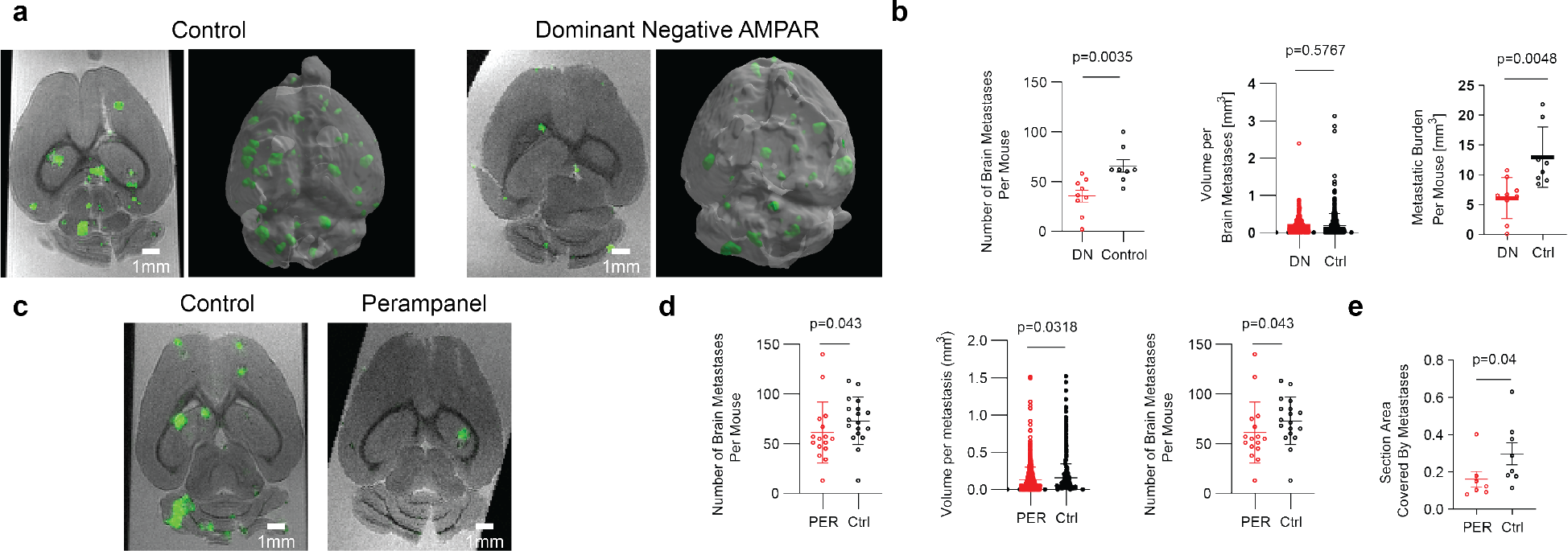
Genetic and pharmacological AMPAR inhibition reduces brain metastatic burden **a**, Representative images and corresponding 3D reconstructions of *ex vivo* T2* MRI of brains from mice injected with Jimt-1 cells expressing control- or GluA2-DN-GFP. Brain metastases are highlighted in green, based on semi-automatic segmentation. **b**, Number of BM per mouse, total metastatic burden and volume per BM for Jimt-1 expressing GluA2-DN-GFP (N=9 mice) vs. control (N=8 mice). P values determined by an unpaired T-test for left and middle graph, and by a Mann-Whitney test for right graph. **c**, Representative *ex vivo* MRI of mice treated with perampanel or control, following intracardial injection with Jimt-1 cells (green, as in **a**). **d**, Metastatic burden, number of BM and volume per BM in perampanel (PER, N=16 mice) treated vs. control (Ctrl, N=18 mice), 28 days following intracardial injection with Jimt-1 tumor cells. P values determined with a Mann-Whitney test for left and middle graph, unpaired T-test for right graph. **e**, Area of A2058 BM on brain sections of mice treated with perampanel (PER, N=10 mice) or control (N=10 mice). P value determined with a Mann-Whitney test.

## Discussion

Here we report that functionally relevant synaptic contacts containing glutamate receptors of the AMPAR subtype exist between neurons and brain metastatic cancer cells of extracranial tumor entities. These findings suggest that such AMPAergic synapses not only support and stimulate cancers of neural origin like gliomas^3–5^, but also brain-seeding breast cancer and melanoma cells, implying a more general mechanism by which particularly malignant tumor entities can hijack neuronal cellular signalling pathways for their own benefit. Genetic or pharmacologic inhibition of these neuron-brain-metastatic synapses resulted in a strongly reduced metastatic burden, hence paving the road for a new, clinically actionable concept of metastasis prevention^11, 27^.

In gliomas, the cell of origin is most likely a neural progenitor or stem cell, such as the oligodendrocyte precursor cell (OPC)^7^. Since it is known that OPCs also receive glutamatergic synaptic input via Ca^2+^- permeable AMPAR^8^, the finding of neurogliomal synapses^3–5^ could have been interpreted as a lineage-restricted recapitulation, or hijacking, of neurodevelopmental features by the primary brain tumor cell. A similar principle could be assumed for extracranial cancers of neuroendocrine origin which also seemingly depend on glutamatergic synaptic signaling^6, 28^. The existence of *bona-fide* neuron-cancer synapses on breast cancer and melanoma cells reported here points towards a much wider pathobiological role of neuron-tumor synapses. It has been shown that even human embryonic kidney cells, when expressing synaptogenic proteins such as neuroligin or synCAMs, attract co-cultured neurons to form synapses^29–31^. Interestingly, while neuron-tumor synapses could not be detected in brain tumor entities of lower malignancy (oligodendrogliomas or meningeomas^3^), a subpopulation of cancer cells in particularly aggressive, incurable primary^3–5^ and also secondary (this manuscript) brain tumor types receives neuronal synaptic input that drives overall disease progression. A key feature of neurogliomal synapses is the expression of Ca^2+^-permeable AMPARs that contribute to a distinct Ca^2+^ signal in the glioma cell^3^. The discovery of such neuronal activity-dependent Ca^2+^ transients in breast cancer and melanoma cells reported here suggests a common mechanism of how neuron-cancer synaptic interactions can be translated into growth-promoting signals.

While previous work reported a proximity mechanism enabling synaptically released glutamate to activate extrasynaptic NMDARs on metastatic breast cancer cells in a pseudo-tripartite configuration^12^, our study reveals direct, *bona fide,* synaptic contacts between neurons and metastatic cancer cells that contain postsynaptic AMPARs. Importantly, these NBMS underly crucial initial steps of metastatic brain seeding and support survival and early proliferation of metastatic cells in the alien microenvironment. In contrast, synaptic proximity signalling appears limited to later stages of macrometastatic growth^12^. The prevention of BM by the clinically approved AMPAR inhibitor perampanel reported here offers a particularly feasible road for clinical translation^13, 24, 26^. Given that more than 90% of cancer patients die of metastasis^32^, it will be of great relevance to test whether similar neuron-cancer synapses exist and can stimulate metastatic organ colonization in extracranial sites as well. After all, glutamatergic synaptic signaling appears to exist in the normal and diseased peripheral nervous system^33, 34^. Remarkably, metastases of various solid and hematological cancer types show a more neuroendocrine-like transcriptomic signature than the primary tumor, resulting in shared vulnerabilities of these neuronal states^35^. Taken together, for the field of brain metastasis, this study demonstrates that direct and indirect synaptic interactions with distinct pro-metastatic effects can collaborate over the course of organ metastasis, providing a roadmap for further studies on how neuron-cancer interactions can govern the metastatic cascade, and how to prevent this. Whatever we find in the future, the discovery of such synaptic contacts on cancer cells from common tumor types that originate outside the nervous system is unexpected and opens a new chapter in the field of Cancer Neuroscience.

## Materials and Methods

### Cancer cell culture

Brain-tropic Jimt-1 (human breast cancer, ER-, PR-, HER2 amplification, trastuzumab resistant, p53-/-, a kind gift from Patricia Steeg, originated from: RRID:CVCL_2077) and A2058 (human melanoma, BRAF- V600E+/-, PTEN+/-, RB1+/-, p53-/-, RRID:CVCL_1059) were cultured in DMEM, 10% fetal bovine serum (FBS) and 1% penicillin/ streptomycin (pen/strep). The mouse brain-tropic mammary adenocarcinoma E0771 cells (triple negative) were a kind gift from Patricia Steeg, and originated from RRID:CVCL_GR23. E0771 were cultured in DMEM with 10% FBS, 1% pen/strep. The patient-derived cell lines from human melanoma BM DDMel31 (*BRAF*-V600E-, *NRAS* Q61+) were cultured in RPMI medium with 10% FBS, 1% penicillin-streptomycin-amphotericin B and 1% L-glutamine. In order to exclude the possibility of establishing tumor associated fibroblasts, the primary cell lines were tested for their *NRAS* or *BRAF* mutation status and for their expression of the melanoma markers Melan A, tyrosinase and HMB45.

All cell lines were stably transduced with lentiviral vectors for imaging purposes. Cytosolic expression of GFP or tdTomato was achieved by transduction with plKO.1-puro-CMV-TurboGFP (SHC003, Sigma-Aldrich, USA) or cytoplasmic tdTomato (LeGo-T2, plasmid #27342, Addgene, USA). For Ca^2+^-imaging, cells were transduced with a Twitch-3A vector (kind gift of Olga Garatschuk, Tübingen, and Oliver Griesbeck, Munich) or pHAGE-RSV-tdTomato-2A-GCaMP6s vector (#80316 Addgene). Both plasmids allow simultaneous Ca^2+^ activity recordings and morphometric analyses of the cancer cells^36^. All non-primary human cell lines were authenticated based on Single Nucleotide Polymorphysm (SNP) typing. Cells were used for experiments between passage two and six after thawing. During the whole duration of the study, all cell lines were tested via PCR every three months for mycoplasma contaminations.

### Neuron-tumor co-cultures

Neuronal co-cultures were performed as described previously^3, 5^. In brief, E19 embryo-derived cells were cultured on 12 mm coverslips, which were situated in 24-well plates pre-coated with poly-L-lysine. These cells were seeded at a concentration of 90,000 cells per square centimeter and sustained in a neurobasal medium from Invitrogen, enhanced with a 2% (v/v) B27 supplement (50x) and 0.5 mM L-glutamine. After a week of in vitro culture (7 DIV), we seeded 1,000 mechanically separated, brain metastatic cells per well for neuron-tumor co-cultures.

### Animals and surgical procedures

All animal procedures were performed in accordance with the institutional laboratory animal research guidelines after approval of the local governmental Animal Care and Use Committee (Regional Council Karlsruhe, Germany, 35-9185.81/G-220/16, 35-9185.81/G-132/16, 35-9185.81/G-50/19, 35-9185.81/G-273/19, 35-9185.81/G-110/21). Efforts were made to minimize animal suffering and to reduce the number of animals used according to the 3R’s principles. All mice were routinely checked for clinical endpoint criteria. Mice (>8 weeks old) were anesthetized with ketamine/xylazine and injected in the left ventricle with 500.000 tumor cells diluted in sterile PBS. Human brain metastatic cells were injected in *Foxn1* Nu/Nu (Charles River, Germany) or *NOD-scid IL2rγ*^null^ (NSG, initially generated by Jackson Laboratory, internal breeding at the German Cancer Research Centre) mice. E0771 cells were intracardially injected into C57BL/6J mice (Janvier Labs). Female mice were used for the breast cancer BM models, males for the melanoma models. The mice brains were isolated and prepared for cryo- or vibratome sectioning. For intravital imaging, the mice received a chronic cranial window with a titanium ring as described previously^13^ at least 3 weeks prior to intracardial injection.

### Intravital two-photon microscopy

Cranial window- and tumor-bearing mice were anesthetized with isoflurane in 100% O_2_. To achieve full narcosis, the mice were first exposed to up to 5% isoflurane and during imaging were kept at ≤1% isoflurane in 100% O_2_. TRITC-Dextran (500 kDa, 52194, Sigma Aldrich, 5 mg mL^-1^) or FITC-Dextran (2000 kDa, FD2000s, Sigma Aldrich, 5 mg mL^-1^) was injected in the tail vein to visualize the brain vasculature. During imaging, the body temperature of the mice was kept at 37 °C using a controllable heating pad. The mice were imaged using a Zeiss multiphoton LSM7 or LSM980 (both Zeiss) equipped with an Discovery NX two-photon tunable femtosecond laser (Coherent). To visualize the fluorophores, either 850 nm (TRITC-dextran) or 950 nm (GFP, tdTomato, GCaMP6s) and a band-pass filter of band pass 500–550 nm/band pass 575–610 nm was used. To visualize TWITCH3A, 860 nm was used to excite CFP (for Förster Resonance Energy Transfer) and a band pass 460–500 nm/band pass 525–560 filter was used to detect CFP and mVenus fluorescence. Gain was set in between 600-850, laser power was kept as low as possible.

### Ca^2+^ imaging

For awake *in vivo* imaging of Ca^2+^ transients in tumor cells, mice were injected cortically, as described previously^36^, with 100.000 brain-tropic A2058 cells transduced with pHAGE-RSV-tdTomato-2A- GCaMP6s. Ca^2+^ imaging was performed during day 14 to day 20 post injection. Mice were immobilized with an implanted cranial ring while remaining mobile within an elevated Mobile HomeCage® (Neurotar, Helsinki). Prior to the in vivo imaging experiment, mice underwent a training regimen of three to four sessions lasting 15 to 60 minutes each to ensure a stress-free environment.

In order to relocate tumor regions between awake recordings and recordings under anaesthesia, 100µl of TRITC-Dextran (see above) was injected before imaging in order to obtain angiograms of the mouse brain. Mice were always imaged awake firstly in order to avoid carryover or long-term effects of anaesthesia affecting subsequent recordings. For each condition, 1000x1024x1024 frames of Ca^2+^ activity were obtained with a frame-time of 2.52s and a pixel size of 0.592. After imaging the awake condition, mice were injected intraperitoneally with a ketamine/xylazine narcosis, retransferred on a heating pad underneath the microscope and the angiogram was used to relocate to the same tumor region. Subsequently, intravital microscopy was repeated with the same settings while adjusting the laser power to achieve similar signal to noise ratios where necessary.

### Ca^2+^ imaging analyses

Images were stabilized in ZEN Black with Timeseries Alignment using the tdTomato (cytoplasmic staining of the tumor cells) channel as reference. Images were then transferred to a custom-written Python pipeline. Briefly, to remove shifts due to movement within a timeseries, the structural similarity (SSIM) between every consecutive image pair *t* and *t+1* was calculated using the implementation in the scikit- image package^37^. If the SSIM for an image-pair was greater than two standard deviations of the mean of all SSIMs of that time series, the *t+1* image was removed from the time series due to large shifts. The remaining frames of each timeseries were then stabilized using the NoRMCorre algorithm^38^ implemented in CaImAn^39^, again using the tdTomato channel as reference. For semi-automatic segmentation, the resulting motion-corrected timeseries of size (*x,y,t*) were then summed up along the temporal axis resulting in a 2D image of the size (*x,y*) and normalized for brightness and contrast using the Contrast Limited Adaptive Histogram Equalization (CLAHE) algorithm implemented in scikit-image^37^. The resulting projected images were then semi-automatically segmented: A custom-trained cellpose^40^ model was applied to the image, the resulting masks of cell bodies were manually inspected for accuracy and undetected cells were manually segmented. With this method, 25% of cells in a FOV could be automatically segmented and the remaining 75% needed to be manually annotated. The label image, consisting of the masks of every cell body, was then applied to the GcAMP6s channel and mean fluorescence intensity per mask over time was extracted using scikit-image^37^. Individual traces of every cell were then detrended using the sinc_filter method from PyBoat^41^, and then the zscore for every time point was calculated. The z score-transformed traces were used for peak detection with the find_peaks function of the scipy python package^42^. For each image pair (awake vs sleep) the same parameters of the peak detection were used. The total number of peaks per timeseries was then normalized to the number of cells and the number of frames to give Ca^2+^ peaks per 100 cells per 10 minutes.

### Multimodal correlative microscopy

Cranial window-bearing mice were imaged using intravital microscopy 3-7 days following intracardial injection with GFP-expressing Jimt-1 cells, as described above. Following identification of tumor cells growing in the perivascular niche and obtaining 3D z-stacks showing both the tumor cells and the vasculature, the mice were sacrificed under deep anesthesia by perfusion fixation. To retrieve the cancer cells for 3D electron microscopy (EM), the protocol was used as described before^19, 43^. In summary, the region of interest was marked post-mortem using near-infrared branding^44^ which enabled dissecting the tissue containing the tumor cells. Following microwave-assisted sample processing involving infiltration of 1% OsO_4_ and 1.5% K_3_Fe(CN)_6_ in 0.1M Cacodylate buffer, 1% OsO_4_ in 0.1M Cacodylate buffer and finally aqueous 1% uranyl acetate, the sample was stepwise dehydrated and embedded in Epon. Small blocks were trimmed around the biopsies and imaged using a x-ray MicroCT to visualize the sample in 3D. Correlation between the intravital microscopy 3D volume and the x-ray volume was performed using Amira (Thermo Fisher Scientific), enabling to accurately map the position of the micrometastases in the resin block. Targeted trimming exposed the region of interest for subsequent serial sectioning and transmission electron microscopy imaging using a CM120 Biotwin (FEI Company, Thermo Fischer Scientific) at 120 kV, using a Keen View CCD Camera (Soft Imaging Solutions, Olympus) or a Tecnai F30 Field Emission Gun at 120-300 kV. TEM images were aligned and segmented using TrakEM (Fiji^45^) or segmented using IMOD^46^ (Boulder Laboratory, University of Colorado).

### Electron microscopy

The procedure for sample preparation was carried out in accordance with previously established methods^47^. Two weeks following intracardial injection, the animals bearing brain metastases were administered a deep anesthesia using a combination of ketamine and xylazine. Subsequently, they were perfused transcardially with a solution containing 4% (w/v) paraformaldehyde (PFA) in 1x phosphate-buffered saline (PBS, Sigma). Upon removal, the brains were postfixed in a 4% PFA solution for a duration of four hours.

Sections of 100-200 µm thickness were then cut using a vibratome (Leica VT1000S). The slices from the xenograft brain were examined under a widefield fluorescence microscope (Leica DM6000) to detect the endogenous fluorescence of brain metastatic cells. For DAB labeling, samples were next immersed in a 10% (w/v) sucrose solution (Sigma) in PBS for a duration of 10 minutes, followed by a 12-15 hour incubation in a 30% sucrose solution. They were then subjected to a freeze-thaw process in liquid nitrogen twice, each for 5 seconds, before being placed in a blocking solution composed of 5% FBS in PBS, and left at room temperature (RT) for 1 hour. The slices were incubated overnight at 4°C with specific antibodies. Alternatively, the samples were processed after dissection at the widefield microscope with a heavy metal stain without DAB labeling and metastatic cells were ultrastructurally identified.

For the DAB precipitate labeling, samples were incubated with the secondary antibody, a biotinylated anti-mouse AB (abcam (ab6788), 1:500, in blocking solution), for 12-15 hours at 4°C. After washing thrice with PBS, the samples were treated with Vectastain ABC-kit (Linaris) solution for 1 hour at RT. The samples were then incubated in a glucose-DAB solution (glucose: 2 mg/ml, DAB: 1.4 mg/ml, dissolved in PBS) for 10 minutes, followed by a 1-hour incubation in a glucose-DAB-glucose oxidase solution (glucose oxidase: 0.1 mg/ml, Serva). This process facilitated the formation of an electron-dense precipitate. The success of the reaction was monitored using widefield light microscopy. All samples were subsequently processed as described previously, embedded in resin and cut with an ultramicrotome (Ultracut S, Leica).

### Acute brain slice preparation

Mice with brain metastases were put under deep anesthesia and then quickly decapitated. Their brains were promptly excised and submerged in an ice-cold slicing solution composed of the following (measured in mM): 125 NaCl, 2.5 KCl, 25 NaHCO3, 1.25 NaH2PO4, 10 glucose, 75 sucrose, 0.5 CaCl2, and 7 MgCl2. The solution was adjusted to approximately 340 mOsm and bubbled with carbogen (a mix of 5% CO2 in O2). Coronal slices of 300 µm thickness were made using a Leica VT1200S vibratome and preserved in the initial slicing solution at 37°C for 30 minutes.

### Electrophysiology

The experiments were conducted utilizing a Leica TCS SP5 System equipped with a 20x water-immersion lens (NA 1.00). Both brain slices and co-cultured coverslips were bathed in carbogen-aerated aCSF at either room temperature or 35°C. Tumor cells were targeted using intrinsic expression of fluorescent proteins in cancer cells. The EPC10 amplifier from HEKA Elektronik, regulated by HEKA’s Patchmaster software, was employed for these recordings. Voltage data were sampled at 100 kHz and filtered at 1 kHz, while current data were sampled at 25 kHz, subjected to a 10 kHz filter, and then Bessel-filtered at 2.9 kHz. Specific traces in figures were filtered at designated frequencies. Patch pipettes were crafted from borosilicate glass tubes (World Precision Instruments) and exhibited resistances between 2 to 7 MΩ when filled with a specialized solution. Voltage clamp mode maintained a holding potential of -85 mV, and no series resistance compensation was applied during recordings. Additional agents were incorporated into the aCSF that superfused the samples: Tetrodotoxin (TTX, Abcam, 1 µM) and CNQX (Abcam, 10µM). All electrophysiological data were subjected to analysis through custom-coded routines using Igor Pro software from Wavemetric Inc.

### Perampanel treatment and evaluation of BM growth

One week before intracardial injection of tumor cells (brain-tropic A2058 and Jimt-1), NSG mice (>8 weeks old) received food pellets containing 320 mg/kg perampanel (Fycompa, Eisai GmbH) *ad libitum* or control food. Depending on their individual response to perampanel, the dose was increased to a maximum of 960 mg/kg. To analyze the effect of perampanel on BM growth, 2-4 weeks following tumor cell injection head-MRI scans were made from the mice. At the end of the experiment, latest on day 28 post-injection, mice were perfused. The brains of the mice injected with Jimt-1 were imaged using *ex vivo* MRI and those of the mice injected with A2058 were cryosectioned and imaged using an Axio Scan.Z1 (Zeiss). The imaging data from the cryosections was semi-automatically quantified as described below.

### MRI imaging

After harvesting of brains, dissected brains were imaged ex vivo on a high field, experimental MRI system (9.4 T, Bruker 9/20 Biospin). After fixation in PFA overnight, brains were transferred to PBS and imaged using a T2* weighted imaging sequence (80µm isotropic resolution). Metastases were delineated based on their hypointense appearance. Metastatic number and volumes were segmented with interactive machine learning^44^.

### Semi-automatic image analysis of brain metastases

MRI imaging data obtained from the T2* weighted imaging sequence were analyzed using the autocontext workflow of ilastik software, as described previously^48^. In parallel, cryosections bearing brain metastases were also processed using ilastik’s autocontext workflow. Subsequent analyses of the processed cryosections were conducted using the Fiji image analysis software using customized macros.

### GluA2Q und Glua2-DN *in vivo* experiments

Brain tropic Jimt-1 were double lentivirally transduced with cytoplasmic tdTomato und GluA2Q-GFP or a dominant negative GluA2-GFP (Glua2-DN GFP). Proliferation rate *in vitro* was assessed using an xCELLigence Real-Time Cell Analyzer (RTCA) system (Roche Diagnostics)^49^. Mice were intracardially injected with either tdTomato/GluA2Q-GFP or tdTomato/Glua2-DN GFP expressing cells. Following perfusion fixation, the brains of the mice were imaged using *ex vivo* MRI.

### RNAseq data analysis

#### Intracohort transcriptomic data pre-processing

Gene expression data of the AMPAR subunit-genes *GRIA1*, *GRIA2, GRIA3* and *GRIA4* in human BM tumor samples and a smaller cohort of primary melanoma and breast cancer tissue samples were obtained from the RNAseq data set of the PreventBM consortium. The use of patient tissue samples for RNAseq in PreventBM was approved by the institutional review board of the Medical Faculty, Heinrich Heine University Düsseldorf (study number: 5717). All patients provided written informed consent for the use of their tissue samples and associated clinical data for research purposes. Paired-end RNA sequencing following library preparation with the lllumina VAHTS total RNAseq Library Prep Kit was performed at the Genomics & Transcriptomics Laboratory, Center for Biolological and Medical Research, Heinrich Heine University Düsseldorf on an lllumina HiSeq 3000 instrument. Alignment to the reference genome was performed using STAR (Version 2.5.3a) algorithm^50^, merging and duplication marking was done with Sambamba (Version 0.6.5)^51^. The output was converted to sorted BAM files withSAMtools (Version 1.6)^52^.

#### TCGA data pre-processing

The R/Bioconductor package TCGAbiolinks^53^ was used to download harmonized transcriptomic data in HTSeq-Counts and HTSeq-FPKM format from three TCGA cohorts: Breast invasive carcinoma (BC), lung adenocarcinoma and skin cutaneous melanoma. ENSEMBL gene identifiers were converted to HGNC symbols using the R package biomaRt^54^. The R package GeoTcgaData was utilized to transform fragments-per-kilobase-million (FPKM) values into TPM values. Gene expression matrices (gene x sample) were generated and counts for all duplicated gene names were replaced with the average. Clinical annotation data was downloaded with TCGAbiolinks.

#### Transcriptomic data analysis

The inhouse PreventBM patient cohort contained RNAseq data from 95 BM samples, 22 primary tumors and 2 normal brain samples. Primary tumor data included (TCGA-BRCA(n=1102), TCGA-LUAD (n=533) and TCGA-Melanoma (n=103) cohorts). For gene expression analysis in both cohorts, matrices (gene x sample) containing raw counts were filtered for at least 5 total reads per gene and then subjected to DESeq2-based normalization^55^. Subsequently, we applied variance stabilizing transformation (VST) to stabilize the variance across the mean and obtain count values that are approximately homoscedastic^56^.

#### AMPA receptor gene expression analysis

To examine transcriptional levels of glutamate receptors of the AMPA subtype, we defined the gene signature for AMPAR by its core components as follows: *GRIA1, GRIA2, GRIA3* and *GRIA4*. DESeq2- normalized and VST-transformed counts from both cohorts were filtered for the GRIA genes and normalized as Z-scores. For each patient, we used the sum of the Z-scores for all *GRIA* genes as the "AMPAR score".

#### GRIA2 editing analysis

*GRIA2* mRNA editing status in the PreventBM cohort was assessed using the mpileup utility from SAMtools^52^, which summarizes the coverage of mapped reads on a reference sequence at single base pair resolution. All reads at chr4:157336723, which is the position where GRIA2 is post-transcriptionally edited (A>G), were counted and we calculated the GRIA2 editing percentage as the number of variant reads (G) divided by the total number of reads. Samples with a total number of mapped reads <5 were excluded from the analysis.

#### Neurotransmitter expression analyses in single cell RNA sequencing data from matched extracranial and intracranial melanoma metastases

Single-cell gene expression data, with the Gene Expression Omnibus accession number GSE185386^25^, was analyzed using Seurat V4^57^. The dataset underwent preprocessing according to the standard procedures available in this R package. Cell-type categorization was conducted using the annotations published alongside GSE185386. All subsequent analyses also utilized Seurat V4^57^.

### Statistical analyses

The results of all quantifications were transferred to the Graphpad Prism (GraphPad Software) to test the statistical significance with the appropriate tests (data were tested for normality using the D’Agostino & Pearson normality). Statistical significance was assessed by the two-sided Student’s t-test for normally distributed data. Otherwise, a Mann–Whitney test was used for non-normal distributions. The data shown in Fig. 4c-h and Supplementary Fig. 3b-g was generated with the open-source statistical programming language R (version 4.0.0).

## Data availability

Single-cell gene expression data have been previously published and can be accessed with the Gene Expression Omnibus accession number GSE185386. Bulk RNA-sequencing data were accessed from the TCGA.

## Code availability

Custom-written code is available upon reasonable request.

## Acknowledgements

The authors would like to thank the CNS Tumor Tissue Bank Düsseldorf, the Biobank of the University Tumor Center (UTZ) / Center for Integrated Oncology Düsseldorf, as well as Iris Helfrich (Ludwig- Maximilians-University, Munich), Katrin Lamszus, Harriet Wikman, Volkmar Müller (all University Hospital Hamburg-Eppendorf, Hamburg), and Anna Sophie Berghoff (Medical University of Vienna, Vienna) for contributing patient samples for the PreventBM cohort. We thank Manuel Fischer (University of Heidelberg) for support of MRI measurements. The authors thank Yvette Dörflinger and Simone Hoppe for help with electron microscopy. We thank Yannick Schwab, Jacky Goetz, Bernhard Ruthensteiner, Luc Mercier and Nicole Schieber for their support with intravital correlative microscopy. The authors thank Carlo Beretta with help in establishing image analysis pipeliens. The authors thank Patricia Steeg for providing the brain-tropic Jimt-1 and E0771 cells. The authors thank Lukas Geckeler and Jonas Henkenjohann for help with analyzing of calcium imaging datasets. We thank Amit Agarwal and Dwight E. Bergles (Heidelberg University) for providing the plasmids encoding the DN-GluA2-GFP and Glua2Q- GFP. The authors thank the Light Microscopy Facility and the Preclinical Research Unit of the DKFZ Heidelberg, and the Advanced Light Microscopy Facility (ALMF) and the Electron Microscopy Core Facility (EMCF) of the EMBL Heidelberg for their technical support and input. Computational support of the Zentrum für Informations- und Medientechnologie, especially the HPC team (High Performance Computing) at the Heinrich-Heine University is acknowledged. The results shown here are in part based upon data generated by the TCGA Research Network: https://www.cancer.gov/tcga. Figure 1a,f,h; Figure 2a; Figure 4a and h; and Supplementary Fig 2a were created with BioRender.com.

M.A. Karreman and D. Westphal were supported by the Bundesministerium für Bildung und Forschung (BMBF) within the framework of the e:Med research and funding concept (01ZX1913A and 01ZX1913D). This study was funded by the Deutsche Krebshilfe (German Cancer Aid), Priority Program "Translational Oncology", #70112507, "PreventBM-Preventive strategies against brain metastases" (addressed to F. Winkler). The work was also supported by the Deutsche Forschungsgemeinschaft (DFG, German Research Foundation), SFB 1389, UNITE Glioblastoma, project ID 404521405 (addressed to V. Venkataramani, M.A. Karreman, M.O. Breckwoldt, W. Wick and F. Winkler), project number VE1373/2-1 (addressed to V. Venkataramani) and project number 259332240/RTG 2099 (addressed to M.A. Karreman and F. Winkler), the Hertie Stiftung (addressed to M.A. Karreman). V.Venkataramani received financial support from the Else Kröner-Fresenius-Stiftung (2020-EKEA.135), Heidelberg University and Research Seed Capital (RiSC) from the Ministry of Science, Research and the Arts Baden Württemberg. We furthermore acknowledge the data storage service SDS@hd, supported by the Ministry of Science, Research and the Arts Baden-Württemberg (MWK) and the DFG through grant INST 35/1314-1 FUGG.

M.O. Breckwoldt was supported by the Emmy Noether program of the German Research Foundation (DFG, BR 6153/1-1).

## Author contributions

F.W., T.K., V.V. and M.K. conceptualised and supervised the study and wrote the manuscript with input from all co-authors. V.V. and M.K. performed the experiments, analysed the data and contributed to all aspects of the study. In particular, V.V. performed the electrophysiology, co-cultures, electron microscopy, expression analyses, and developed image analyses pipelines. M.K. performed calcium imaging *in vivo*, longitudinal intravital microscopy, correlative microscopy, expression analyses, intracardiac injection and other *in vivo* experiments. NBMS were first discovered by V.V. on ultrastructural BM data from M.K. L.N. performed electrophysiological recordings *ex vivo* and on co-culture. N.H. performed intravital microscopy, tumor cell injection, and *in vivo* calcium imaging. N.H. also developed an automatic data analyses pipeline and analysed the data. C.T. performed *in vivo* experiments and intravital microscopy. C.M. performed transductions and *in vitro* experiments. L.M., S.M. (supervised by B.B.) and

M.S. performed bioinformatics analyses of RNA expression data. G.R., J.F. and K.K. were responsible for the histological classification, nucleic acid extraction, clinical data annotation and RNA sequencing of the PreventBM cohort. D.W. provided patient-derived melanoma BM cell lines and input. M.B. performed MRI imaging. W.W. provided conceptual input and supported the project.

---

**Supplementary Figure 1:**
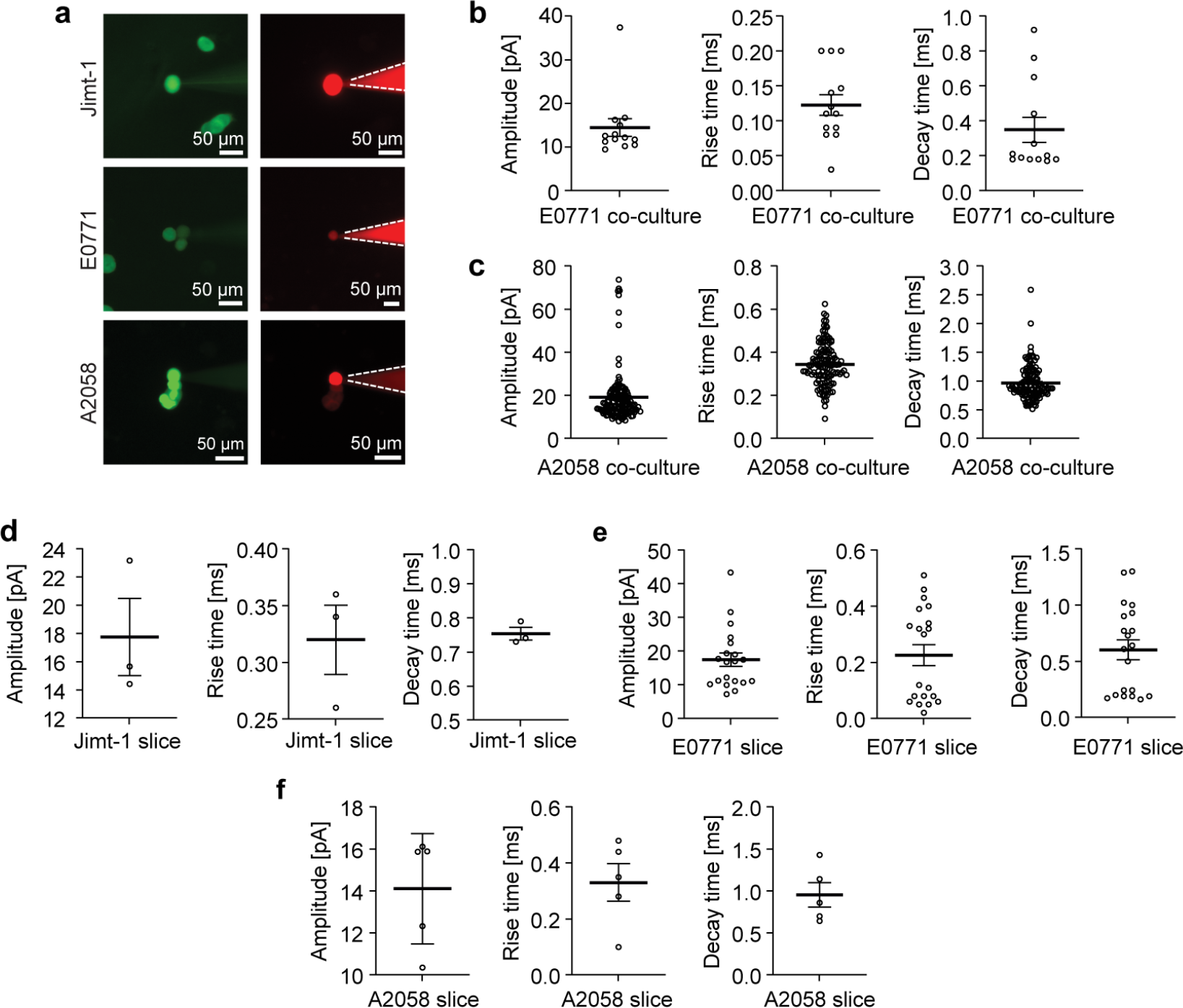
Electrophysiological characterization of EPSCs in breast cancer and melanoma brain metastases models *in vitro* and *ex vivo* **a**, Whole-cell recording from Jimt-1 (human breast cancer), E0771 (mouse mammary cancer) and A2058 (human melanoma) brain metastatic cells stably expressing GFP (left panels) and injected with red fluorescent dye through the patch pipette (right panels). **b**-**f,** Amplitude, rise- and decay time of EPSCs *in vitro* co-cultures of neurons and E0771 (**b**) and A2058 (**c**), and in acute brain slices of mice intracardially injected with Jimt-1 (**d**, 14 days post-injection), E0771 (**e**,10-14 days post-injection) and A2058 (**f**, 14 days post-injection).

**Supplementary Figure 2:**
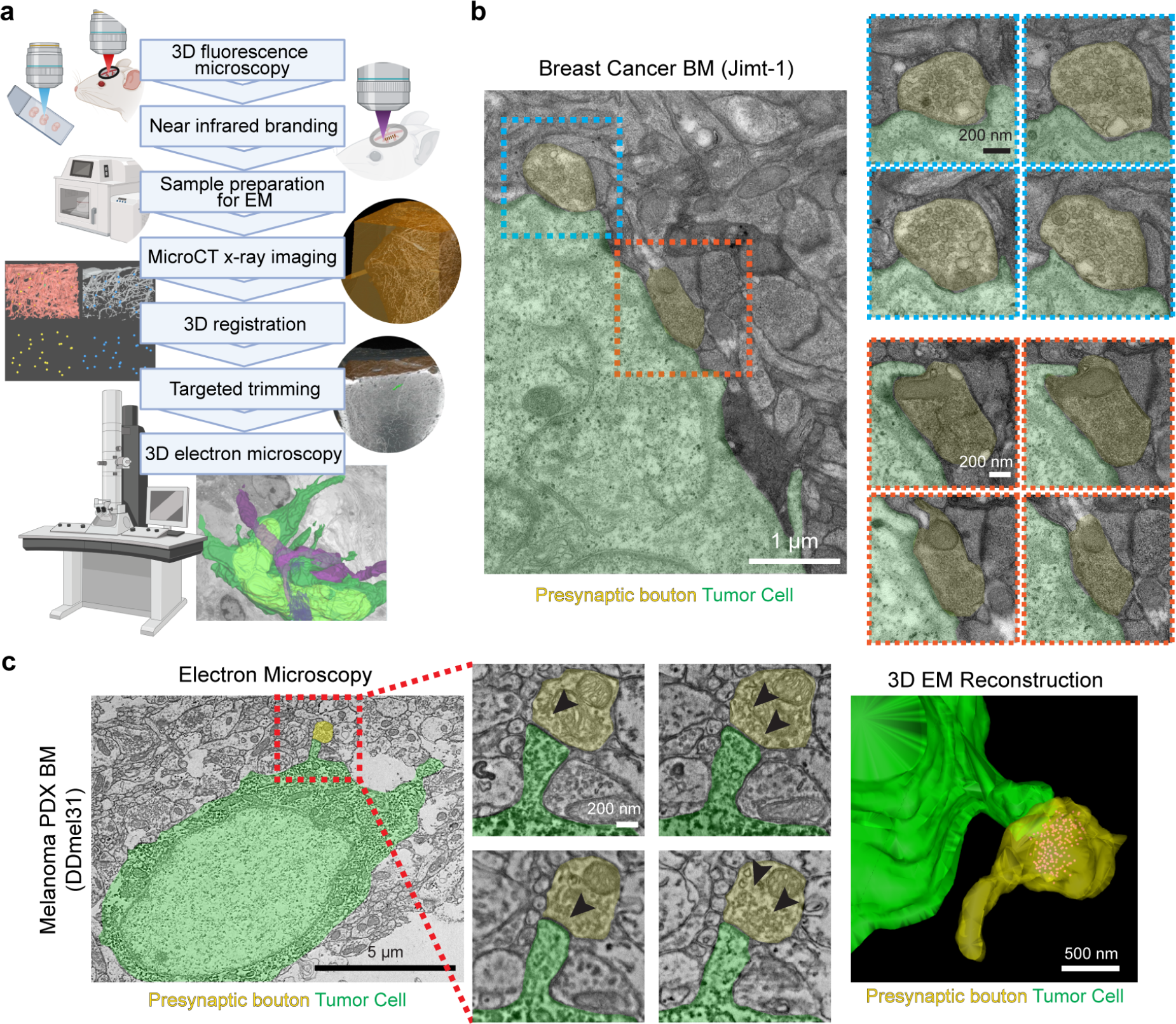
Neuron-BM synapses in preclinical models and patient resectates **a**, Schematic overview of the correlative multimodal microscopy workflow used to visualise early, perivascular brain metastases. **b**, Representative micrographs of neuron-brain metastases synaptic structures at Jimt-1 BCBM, 4 days following intracardial injection. **c**, Immuno-electron microscopy of DAB (3,3′-Diaminobenzidine)-marked DDMel31 melanoma PDX brain micrometastasis. DAB can be recognised as electron-dense deposits in the tumor cell cytoplasm. Melanoma BM is marked in green, presynaptic bouton in yellow. Arrowheads show presynaptic vesicles. The 3D EM reconstruction shows the presynaptic vesicles (red).

**Supplementary Figure 3:**
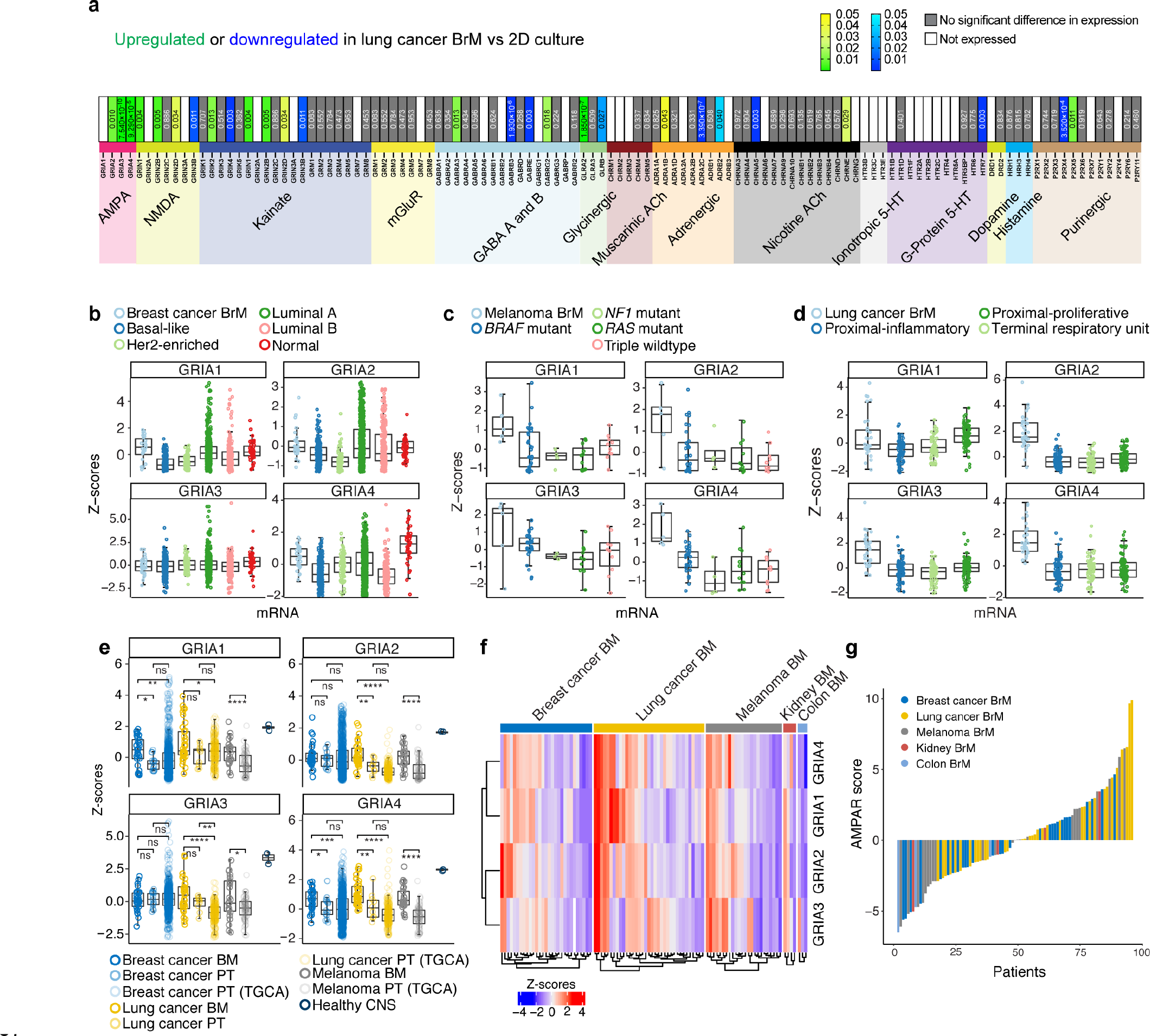
Expression of neurotransmitter receptor genes in BM and primary tumors of various human tumors of melanoma, breast- and lung cancer **a**, Differential regulation of neurotransmitter receptor genes (indicated below) in human lung cancer cell cells (H2030-BM3) grown in 2D culture vs. in the forebrain of mice^18^. Significant upregulation is indicated in yellow/green, downregulation in light to dark blue. Grey indicates no significant difference in RNA expression, white cells indicate no expression. AMPAR genes *GRIA2-4* are significantly upregulated in LUAD-BM cells grown in the brain compared to 2D culture. **b-d**, Boxplots of the expression of *GRIA1-4* in brain metastases (PreventBM) and different subtypes of primary breast cancer (**b**), melanoma (**c**) and lung adenocarcinoma (**d**). Expression data on primary tumors from the PreventBM dataset and the Cancer Genome Atlas (TCGA). **e**. Boxplot comparison of z-scored mRNA expression of AMPAR-genes *GRIA1-4* between BM, primary tumor and healthy CNS (n=2) datasets. BC: breast cancer, LUAD: lung adenocarcinoma; Mel: Melanoma; BM: brain metastases; Prim Tumor: primary tumor; CNS: non-pathological central nervous system tissue. P values were determined by an unpaired two-sided Wilcoxon test. **f**. Clustered heatmap of z-scored mRNA expression of *GRIA1-4* in 95 patients with BM from different entities (BC: breast cancer; LUAD: lung adenocarcinoma; Mel: melanoma). **g**. Waterfall-plot for AMPAR scores per patient (as in f), ranked in increasing order from left-to-right.

## Supplementary Videos

**Supplementary Video 1:** Intravital two-photon microscopy showing a time series video recording of Ca^2+^ activity in early Jimt-1 breast cancer brain metastases expressing the fluorescent calcium sensor TWITCH3A. Spontaneous calcium transients in brain metastatic breast cancer cells can be observed.

**Supplementary Video 2:** Intravital two-photon microscopy showing a time series video recording ofI Ca^2+^ activity in early A2058 melanoma brain metastases expressing fluorescent calcium sensor GCaMP6f. Spontaneous calcium transients in brain metastatic melanoma cells can be observed.

**Supplementary Video 3:** Intravital two-photon microscopy of melanoma brain metastases. Time series video recording of of Ca^2+^ activity in A2058 melanoma brain metastases in an awake mouse, followed by Ca^2+^ imaging of the same region while the mouse is anaesthetized. A clear reduction of calcium transients in cancer cells can be observed in mice under anaesthesia as compared to awake mice.

**Supplementary Video 4:** Serial section 3D transmission electron microscopy of NBMS in early Jimt-1 breast cancer brain metastases. First, the raw serial section electron microscopy data are shown, followed by the three-dimensional segmentation and rendering of NBMS. Tumor cells were identified by correlative intravital two-photon and electron microscopy. In the segmentation, the tumor cell membrane is colored in green. The neuronal presynaptic boutons (yellow with red vesicles and blue with red vesicles) can be identified by the rendered synaptic vesicles. Scale bar represents 1 µm.

**Supplementary Video 5:** Serial section 3D scanning electron microscopy of NBMS in early A2058 melanoma brain metastases First, the raw serial section electron microscopy data are shown, followed by the three-dimensional segmentation and rendering of NBMS. Tumor cells can be identified by dark DAB precipitation. In the segmentation, the tumor cell membrane is colored in green. The neuronal presynaptic boutons are depicted in yellow with red presynaptic vesicles. Scale bar represents 1 µm.

**Supplementary Video 6:** Serial section 3D scanning electron microscopy of NBMS in early DDMel31 melanoma brain metastases. First, the raw serial section electron microscopy data are shown, followed by the three-dimensional segmentation and rendering of NBMS. Tumor cells can be identified by dark DAB precipitation, tumor cell membrane is segmented in green. The neuronal presynaptic bouton is depicted in yellow with red presynaptic vesicles. Scale bar represents 1 µm.

